# The Regulation of COX-2, ABCA1 and ABCG1 by the lncRNA PACERR links the inflammatory response and cholesterol homeostasis

**DOI:** 10.1101/2025.06.02.657432

**Authors:** Elizabeth J. Hennessy, Takae Tanosaki, Ronan Lordan, Soon Yew Tang, Nadim El Jamal, Vladimir Shuvaev, Soumita Ghosh, Ujjalkumar S. Das, Robin Joshi, Hu Meng, John N. Snouwaert, Hanna Winter, Lars Maegdefessel, Beverly H. Koller, Garret A. FitzGerald

## Abstract

Cyclooxygenase-2 (COX-2) has an established role in inflammation and its inhibition by non-steroidal anti-inflammatory drugs (NSAIDs) is clinically efficacious^1,2^. However, placebo-controlled trials of COX-2 inhibitors have revealed a cardiovascular hazard. This has been attributed to suppression of COX-2 dependent cardioprotective prostaglandins (PGs), especially prostacyclin, produced by endothelial (EC) and vascular smooth muscle cells (vSMC) ^1-3^. Upstream of these effects, we have found that the COX-2 antisense long non-coding RNA (lncRNA) PACERR interacts with discrete RNA binding proteins to mediate the cell specific COX-2 response to both anti-[high-density lipoprotein (HDL)] and pro-inflammatory [lipopolysaccharide (LPS)] stimuli. PACERR exerts its action both in *cis* and in *trans,* controlling the expression of COX-2 and the cholesterol transport genes, ABCA1 and ABCG1. When cells are exposed to HDL, PACERR is transcribed, leading to the release of bound histone acetyltransferase p300 and its relocation to the promoter of COX-2 which is then transcribed leading to PG formation. In concert, PACERR also releases heterogeneous nuclear ribonucleoprotein L (hnRNPL) which relocates to the ABCA1 and ABCG1 gene loci, controlling mRNA alternative splicing and subsequent cholesterol efflux to HDL. Silencing PACERR, with an antisense oligonucleotide (ASO), augments the COX-2 and cholesterol efflux in response to HDL. In response to HDL, PACERR associates with its neighboring gene, cytosolic phospholipase A2 (cPLA2/PLA2G4A) both interacting with an enhancer element affecting cPLA2 transcriptional activation and escorting the cPLA2 protein to the nuclear membrane. There, it releases arachidonic acid, further activating COX-2 but limiting ABCA1 and ABCG1 by suppressing the formation of the LXR/RXR heterodimers required for their transcription when cholesterol efflux is no longer required^4^. In response to LPS, PACERR controls the action of the NFκB repressive protein, IκBζ. When PACERR is silenced in a humanized mouse model with the ASO, IκBζ is free to bind to the NFκB site in the COX-2 promoter, suppressing transcription, PG formation and systemic inflammation. In summary, these studies reveal dual and complementary mechanisms by which PACERR modulates PG production by COX-2. In response to HDL, it enhances production of prostacyclin whereas it dampens production of the PG response to LPS.

## Introduction

Non-steroidal anti-inflammatory drugs (NSAIDs) are amongst the most used drugs by adults, offering efficient and effective pain relief through the suppression of cyclooxygenase activity and subsequent prostaglandin formation. However, their use is complicated by adverse effects such as the development of gastrointestinal lesions and cardiovascular complications including myocardial infarction, hypertension, stroke and heart failure. Upon an inflammatory insult, COX-2 converts arachidonic acid, a polyunsaturated fatty acid present in the phospholipids of cell membranes, into prostaglandins (PGs), including PGE^2^ which promotes inflammation, proliferation, and angiogenesis and prostacyclin (PGI^2^) which plays a protective role in the vasculature acting as an inhibitor of macrophage cell activation and adhesion while also preventing platelet aggregation, and vascular smooth muscle cell (vSMC) contraction, migration, and growth^5^ ^6,7^. Augmented PGI^2^ is a homeostatic response to limit the consequences of platelet activation *in vivo* ^8^. While suppression of pro-inflammatory COX-2 dependent PGs contributes to NSAID efficacy, inhibition of cardioprotective PGs, such as PGI^2^ largely explain the cardiovascular risk associated with NSAIDs^9^. It is unknown whether the regulated expression of COX-2 contributes to the therapeutic index of NSAIDs.

Atherosclerosis is a chronic inflammatory disease in which lipid rich plaques accumulate along disturbed arterial walls, disrupting blood flow. Global post-natal deletion of Cox-2, as well as endothelial cell (EC) or vSMC specific Cox-2 deletion accelerates atherogenesis, whereas myeloid Cox-2 deletion mitigates progression of the disease in mice ^10-12^. Epidemiological studies have reported an inverse relationship between circulating high-density lipoprotein (HDL) and cardiovascular outcomes, however Mendelian genetics have failed to support mitigation of cardiovascular risk by HDL ^13,14^. Amongst the mechanisms by which HDL may influence cardiovascular risk is through its effects on cholesterol metabolism and inflammation. *In vitro*, HDL upregulates COX-2 and subsequent PGI^2^ formation through CREB (cAMP response element-binding protein) activation^15-19^. The importance of this mechanism *in vivo* is unknown.

Long non-coding RNAs (lncRNAs) hold therapeutic promise due to their specific and regulated patterns of expression in cells and tissues, functioning as signals, decoys, guides or scaffolds ^27-34^. They have no strict sequence conservation restraints like protein-coding genes, and they evolve quite rapidly, so their sequences are often poorly conserved between species. Their interactions with other molecules can result in cellular epigenetic modifications such as changes to DNA methylation status and modifications to histones as well as the remodeling of chromatin, which culminate in changes in the expression of target genes. Although several thousand lncRNAs have been identified in the genome^20-23^, the function of only a limited number has thus far been described^24,25^.

We therefore investigated the interface between the response of COX-2 to HDL and inflammation in the vasculature by identifying lncRNAs mediating the responses. We have uncovered a dual regulatory role for the lncRNA PACERR (<u>P</u>TGS2 <u>A</u>ntisense <u>C</u>omplex-Mediated <u>E</u>xpression <u>R</u>egulator <u>R</u>NA) on the expression and function of COX-2 ^26^. Depending on the stimulus and the cell type expressing PACERR and COX-2, its PG products vary and can promote or restrain inflammation and cholesterol homeostasis. We have found that PACERR expression, like COX-2 is induced by both lipopolysaccharide (LPS) and HDL in vascular cells, including macrophages, vSMCs and ECs. It is the first lncRNA to be implicated in the regulation of PG production.

PACERR exhibits functional orthology in mouse and human; however, the transcripts share little sequence homology, only a single NFκB binding motif. Because of genomic differences seen in the loci containing the mouse and human PACERR genes, we have generated a novel mouse model where the Cox-2/Pacerr (Ptgs2os) locus has been replaced with the human COX-2/PACERR locus to interrogate the role of PACERR in cholesterol metabolism and the acute inflammatory response. We found that when PACERR is silenced with an antisense oligonucleotide (ASO), COX-2 expression and PG production are augmented in response to HDL but are attenuated in response to LPS. Through its interactions with various RNA binding proteins including p300 and heterogeneous nuclear ribonucleoprotein L (hnRNPL), we have identified PACERR as having a key role in the *cis* regulation of COX-2 and the *trans* regulation of the cholesterol transporters ABCA1 and ABCG1 maintaining intracellular cholesterol homeostasis and inflammation thus attenuating cardiovascular risk.

Here, we reveal a discriminant impact of human PACERR in modulating the expression of COX-2 by divergent stimuli regulating both PG formation and cholesterol homeostasis and potentially contributing to the cardiotoxicity of NSAIDs observed in clinical trials.

## Results

### PACERR is expressed in carotid plaques and is induced by HDL dependent on the CREB transcription factor

Previous studies have implicated COX-2 in the progression of atherosclerosis, so we first determined if PACERR is differentially expressed in early stage versus late or advanced plaques from human carotid endarterectomy samples ^27,28^. We found augmented PACERR expression in samples taken from early plaque regions compared to more advanced plaques implicating PACERR in the initiation of early inflammatory events in the vasculature (Figure 1A). Upon examining single cell RNA-sequencing from human carotid atherosclerotic samples, PACERR was most highly expressed in ECs followed by inflammatory macrophages further associating PACERR in early plaque development and involvement in the response to EC dysfunction (Figure 1B). Because of the observed expression of PACERR in human carotid plaque and inflammatory macrophages, we used the human THP-1 monocyte cell line differentiated into macrophages and treated the cells with various ligands involved in cholesterol metabolism and inflammation. Krawczyk *et al* showed that like COX-2, the lncRNA PACERR is responsive to LPS in differentiated U937 macrophages^26^. We found that both PACERR and COX-2 are concomitantly induced by the HDL precursor apolipoprotein A1 (ApoA1), HDL, SR-BI dependent acetylated low-density lipoprotein (acLDL), CD36 dependent oxidized LDL (oxLDL) and LPS (Figure 1C). The transcription start site of PACERR is 364 basepairs from the COX-2 transcription start site, so we anticipated co-regulation of the two genes and that PACERR would be dependent on COX-2. Using the COX-2 selective NSAID celecoxib to inhibit COX-2 enzymatic activity and subsequent PG synthesis, we found that when cells were pre-treated with celecoxib before HDL or LPS, PACERR is induced to the same degree (Figure 1D). Interestingly, we found that COX-2 expression is attenuated when cells were treated with celecoxib before LPS signifying a feedback loop where the suppression of PGs affects COX-2 expression.

**Figure 1.**
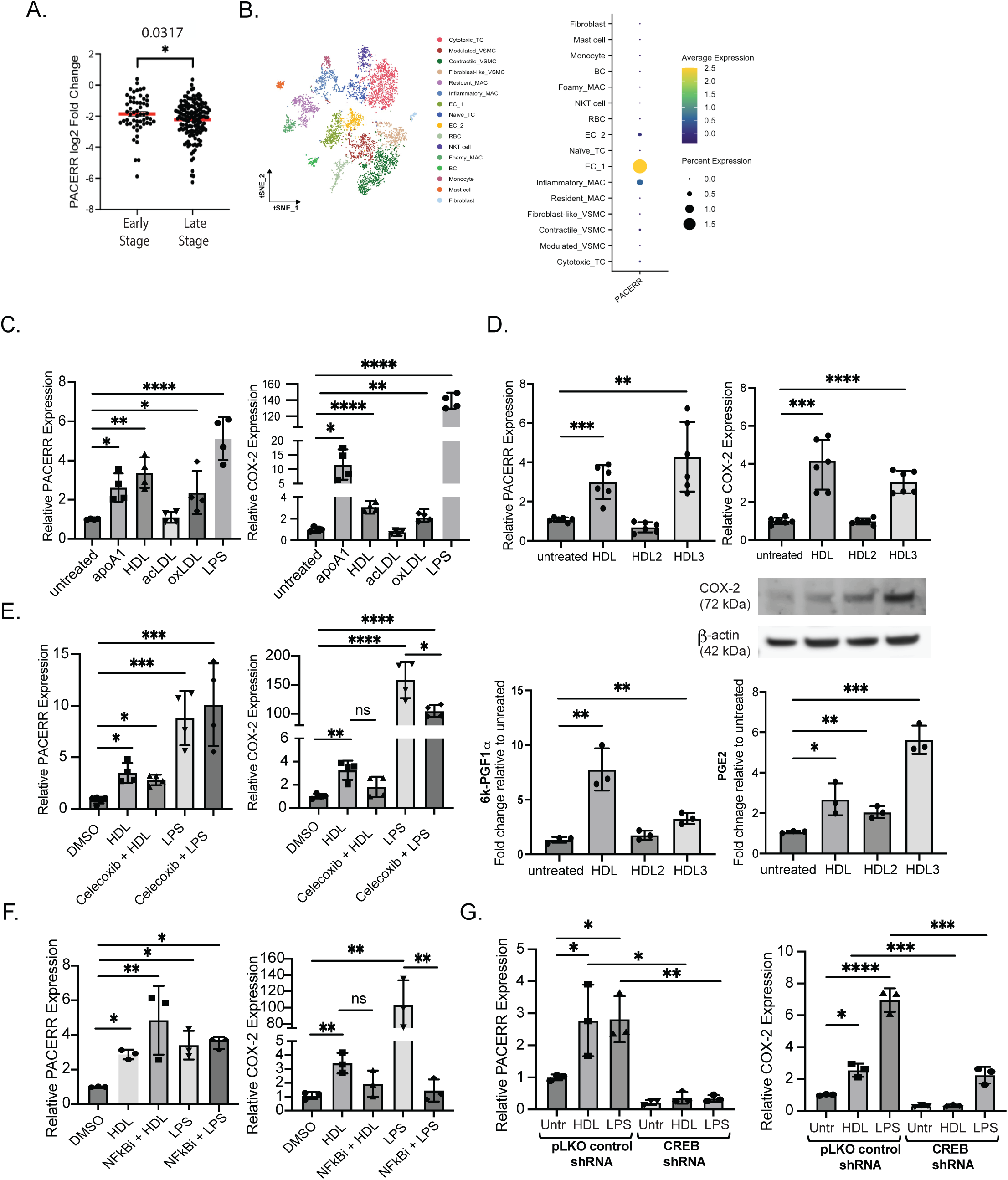
The COX-2 antisense long non-coding RNA PACERR is expressed in carotid plaques and is induced in macrophages in response to HDL. (A) PACERR expression in total RNA isolated from human carotid endarterectomy samples where advanced/central plaques are separated from peripheral plaque and sequencing was done on both. (B) scRNA-seq of carotid plaque samples. (C) THP-1 cells differentiated into macrophages using 10 nM PMA for 48–72 h then treated with ApoA1 (50µg/ml), HDL (50µg/ml), acLDL (37.5µg/ml), oxLDL (10µg/ml), or LPS (100ng/ml) for 24 hours followed by qPCR for gene expression analysis. (D) THP-1 macrophages were treated with 50µg/ml HDL, HDL2 or HDL3 for 24 hours and qPCR for gene expression was used, western blot to measure COX-2 protein levels and mass spectrometry was used to measure 6k-PGF1a/PGI2 and PGE2 in the cell media. (E) THP-1 macrophages were pre-treated with 5µM celecoxib for 1 hour before addition of 50µg/ml HDL or 100ng/ml LPS for 24 hours. (F) THP-1 macrophages were pre-treated with 5µM Bay-117082 for 1 hour before addition of 50µg/ml HDL or 100ng/ml LPS for 24 hours. (G) THP-1 macrophages stably expressing CREB shRNA were treated with 50 µg/ml or 100ng/ml LPS for 24 hours then qPCR was used to measure gene expression. Values are mean ±SEM of three independent experiments. *p<0.05, **p<0.01, ***p<0.005, ****p<0.001 versus untreated.

The COX-2 promoter region is rich with transcription factor binding sites, including multiple NFκB and CREB sites, each controlling its expression and activity in disease states from cancer to inflammation, we anticipated a shared transcription factor paradigm between the two genes^29^. LPS induces COX-2 expression via binding of the NFκB subunits p65 and p50 to transcription factor sites in its promoter region^30^. The transcription of COX-2 is followed by production of pro-inflammatory PGs together with cytokines ^31-34^. Previous studies have implicated LPS induced PACERR in the regulation of COX-2 transcription by sequestering the repressive NFκB p50/p50 homodimer from binding to the NFκB sites allowing for binding of the activating p65/p50 heterodimer and the histone acetyl transferase p300 to be recruited resulting in a more permissive chromatin state^26,35^. We wanted to determine what factors were transcriptionally regulating PACERR. THP-1 macrophages were pre-treated with the NFκB inhibitor BAY-117082 before LPS stimulation^36^, and we found that LPS-induced COX-2 expression was reduced whereas PACERR levels remained elevated following LPS, whereas COX-2 expression was reduced suggesting an NFκB independent mechanism of PACERR induction (Figure 1E). We observed a similar result in THP-1 macrophages pretreated with the NFκB inhibitor BAY117082 prior to HDL where we saw a modest dependency of COX-2 expression on NFκB, but PACERR expression remained comparable in cells pretreated with the inhibitor compared to those treated with HDL alone.

Several studies have shown that the lipoproteins ApoA1 and HDL induce COX-2 expression through CREB signaling independent of NFκB ^32,37,38^, we next determined if HDL also induces PACERR via CREB signaling. We designed shRNAs targeting CREB for stable knockdown of the transcription factor in THP-1 cells. We differentiated these cells into macrophages and treated them with HDL and found a dependency on CREB signaling for both COX-2 and PACERR expression (Figure 1F). Previous studies have shown that CREB promotes an anti-inflammatory immune response through the inhibition of NFκB while also inducing IL-10^16^. Thus, we hypothesize that CREB dependent PACERR induction by HDL is inhibiting NFκB activation limiting inflammation and potentially contributing to the cardio-protective role of COX-2 in the vasculature.

### PACERR regulates COX-2 expression and HDL induced PGs

Because of the observed expression of PACERR in ECs and inflammatory macrophages in human carotid plaques, and it is most highly expressed in arterial tissues in the Genotype Tissue Expression (GTEx) database, we next wanted to see if its expression is differentially modulated in cells of the vasculature including primary peripheral blood mononuclear cells differentiated into macrophages, vSMCs and ECs. We found that HDL induces PACERR and COX-2 in PBMC macrophages, vSMCs and when ECs are exposed to laminar flow of 10 dyn/cm^2^ (Figure 2A-C). Silencing PACERR using an antisense oligonucleotide (ASO) further augmented HDL induced COX-2 expression in the three cell types and the release of PGI^2^ and PGE2 by macrophages and SMCs but PGE2 was suppressed by HDL treatment of ECs, whereas knockdown of PACERR increased PGE2 levels consistent with the increase in COX-2 mRNA expression and protein (Figure 2D).

**Figure 2.**
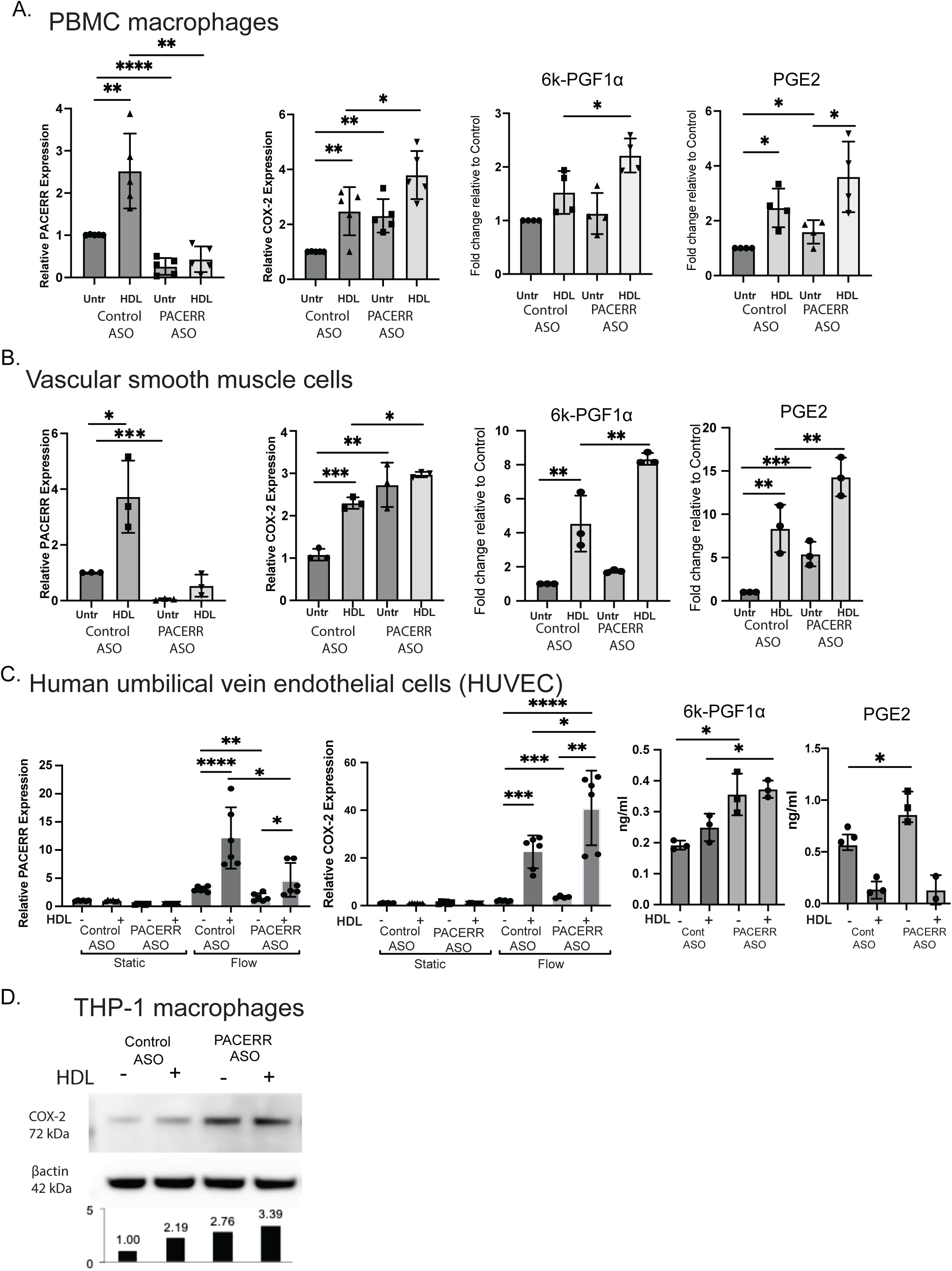
Silencing PACERR with antisense oligonucleotides increases HDL induced COX-2 expression and prostaglandins. (A-C) PBMC macrophages, vSMCs and HUVECs were transfected with 50nm ASO and then treated with 50µg/ml HDL for 24 hours, RNA was extracted and gene expression measured using qPCR. 6k-PGF1α and PGE2 were measured in the supernatant and normalized to total RNA concentration and normalized to untreated cells. HUVECs were subjected to laminar flow conditions 10 dyn/cm^2^ for 24 hours before collection for qPCR. (D) Western blot for COX-2 in THP-macrophages transfected with 50nm control or PACERR ASO and unstimulated or treated with 50µg/ml HDL for 24 hours. Values are mean ±SEM of three independent experiments. *p<0.05, **p<0.01, ***p<0.005, ****p<0.001 versus control untreated.

### PACERR impacts cholesterol metabolism gene expression and cholesterol efflux

Because of the response by macrophages, vSMCs and ECs to extracellular HDL, we next wanted to see if PACERR plays a role in the removal of excess cholesterol from macrophages via HDL, an essential process in the maintenance of cholesterol homeostasis in the body^39^. Human THP-1 macrophages transfected with an ASO targeting PACERR exhibited increased cholesterol efflux (Figure 3A) along with the augmented expression of LXR responsive genes involved in the clearance of excess cholesterol, ABCA1 and ABCG1 (Figure 3B). Interestingly, we also found that PACERR expression is induced in response to treatment of THP-1 macrophages with the LXR agonist T0901317 (data not shown) further signifying a potential role in the regulation of the LXR responsive genes ABCA1 and ABCG1.

**Figure 3.**
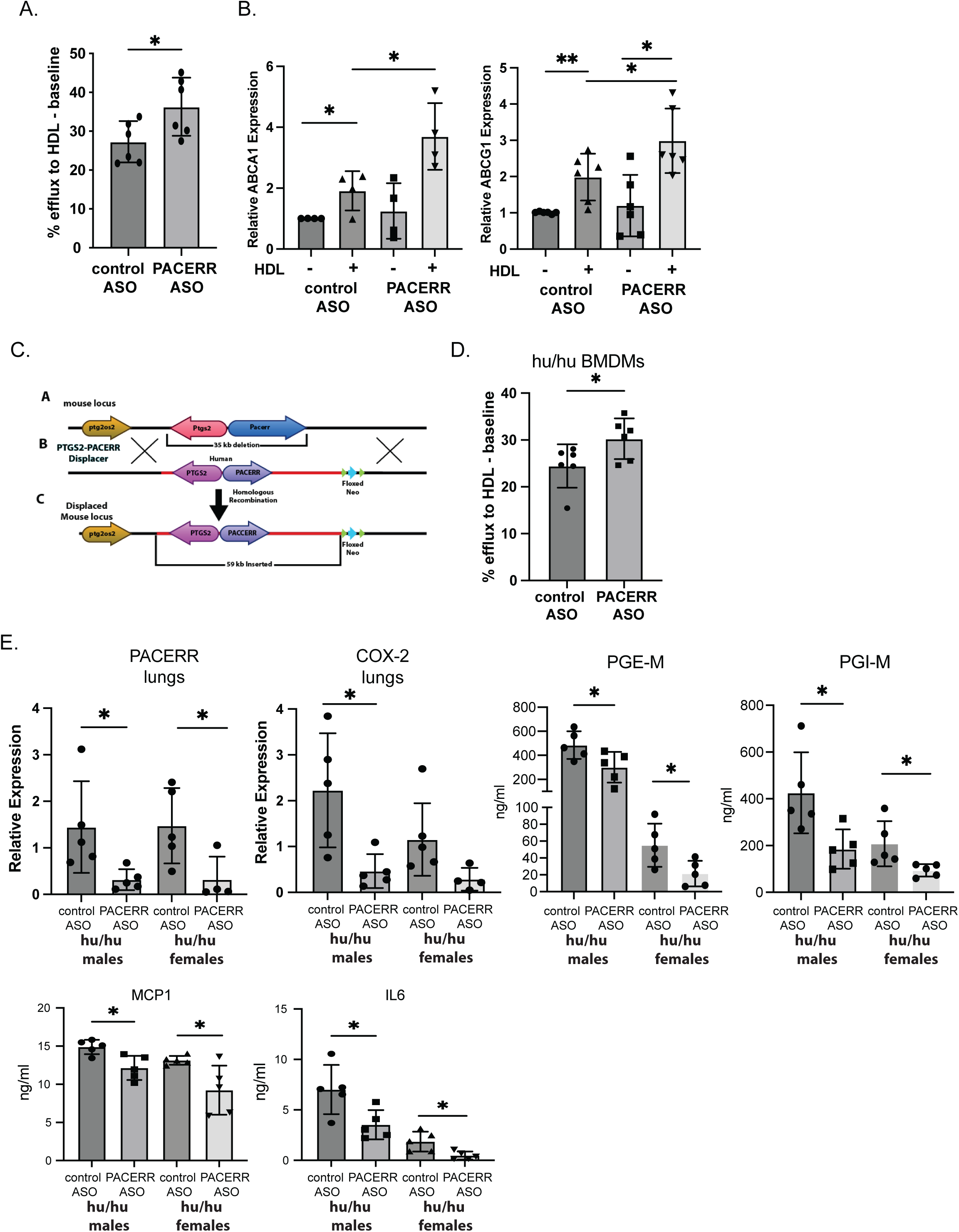
Knockdown of PACERR augments cholesterol efflux to HDL and attenuates the inflammatory response to LPS in a humanized mouse model. (A) THP-1 macrophages were transfected with 50nm control or PACERR ASOs and then loaded with ^3^H-cholesterol and 37.5µg/ml acLDL followed by HDL for efflux of cholesterol. Percent efflux is expressed as a percentage of total cell ^3^H-cholesterol content minus baseline efflux. (B) ABCA1 and ABCG1 expression was then measured in THP-1 macrophages transfected with control or PACERR ASOs and treated with 50µg/ml HDL. (C) Schematic showing the strategy used to generate humanized mice, 35kb of the mouse locus containing Cox-2/Pacerr was replaced with a 59kb syntenic region of the human COX-2/PACERR locus. (D) Cholesterol efflux by BMDMs isolated from the humanized mice (hu/hu) and transfected with control or PACERR ASOs was measured following loading with ^3^H-cholesterol and 37.5µg/ml acLDL followed by HDL for efflux of cholesterol. Percent efflux is expressed as a percentage of total cell ^3^H-cholesterol content minus baseline efflux. (D) Hu/hu mice were retro orbitally injected with PECAM-LNPs encapsulating control ASO or PACERR ASO then intraperitoneally injected with 1µg/kg LPS for 6 hours (n=6 per group). Lungs were isolated and RNA extracted for gene expression analysis. Values are mean ±SEM of three independent experiments. *p<0.05, **p<0.01, ***p<0.005, ****p<0.001 versus control.

The human and mouse genomic regions PACERR exhibit little sequence conservation but share syntenic regions. There is evidence for transcription of the *PACERR* gene in mouse and human. The mouse and human transcripts are both located on chromosome 1 and overlap the promoter region for *COX-2*. The mouse genomic region containing COX-2 and PACERR is ∼25,000 base pairs long. There are two annotated mouse Pacerr isoforms consisting of either 5 exons (∼3000 base pairs) or 3 exons (∼1800 base pairs). The human *PACERR* gene is a single exon that is 825 nucleotides in length. A BLAST search revealed that the mouse and human gene regions share only a 16-nucleotide sequence representing an NFκB site in the *COX-2* promoter region.

In order to study human PACERR *in vivo*, we generated a humanized mouse where 35 kilobases (kb) of the mouse locus containing Cox-2/Pacerr was replaced with a 59kb syntenic region of the human COX-2/PACERR locus (Figure 3C). Our ability to successfully replace the mouse Cox-2 locus with the human COX-2 locus addresses several important points. We were able to humanize the locus despite its complex structure, the high amount of repeat DNA as well as the size of the region. We first isolated bone marrow derived macrophages (BMDMs) from the humanized mice (hu/hu BMDMs) to determine if macrophages from the humanized mouse behaved the same as the THP-1 macrophage cell line transfected with PACERR ASO. We confirmed that hu/hu BMDMs transfected with PACERR exhibited increased efflux of cholesterol to extracellular HDL (Figure 3D).

Because of the induction of PACERR in response to LPS *in vitro*, we next wanted to see how the humanized mice would respond to acute inflammation following intraperitoneal injection of LPS. Lungs harvested from humanized mice pre-treated with PECAM labeled LNPs encapsulating PACERR ASO displayed reduced PACERR and COX-2 expression, PGE-M and PGI-M urinary metabolites as well as attenuated systemic inflammation as seen by circulating plasma IL-6 and MCP1 (Figure 3E). This is in agreement with the previous study implicating PACERR in the COX-2 response to LPS where decreased PACERR led to enhanced binding of the p50 subunit to the COX-2 promoter and prevention of COX-2 transcription^26^. These results signify that silencing PACERR with an ASO can result in a beneficial increase in the removal of excess cholesterol from macrophages as well as reduce systemic inflammation.

### p300, IkBζ and hnRNPL bind to PACERR RNA to mediate effects on COX-2, ABCA1 and ABCG1

Many lncRNAs regulate the expression of genes and pathways through their interactions with proteins, such as transcription factors, ribonucleoproteins and chromatin modifying molecules. To identify RNA binding protein (RBP) partners of PACERR we used an unbiased mass spectrometry proteomic screen and confirmed potential interacting partners using western blotting with antibodies for the specific proteins identified^40^. We focused on proteins relevant to this study involving cholesterol metabolism and inflammation. PACERR bound to p300 (CREB binding protein), IkBζ and_the heterogeneous nuclear ribonucleoprotein L (hnRNPL) (Figure 4A). p300 acts as a transcriptional coactivator by binding to phosphorylated CREB which then binds to cAMP response elements (CREs) to regulate gene expression. p300 functions as a histone acetyltransferase to modify histones by adding acetyl groups and making chromatin more accessible. Because of the CREB dependency for PACERR and COX-2, we examined the transcriptional regulation of COX-2 by PACERR via its interaction with p300. The mass spectrometry screen did not identify the previously implicated protein target p50 but did identify IκBζ ^26^. Atypical IκBs (e.g. BCL3, IκBζ, IκBNS) are mostly inducible and mainly localized to the nucleus. They act as repressing co-factors of NFκB-mediated target gene expression. The interaction of IκBζ with p65/p50 suppresses gene expression by recruiting HDAC proteins to promoters^41^. We identified a basal interaction with IkBζ which preferentially associates with p50 versus p65 to regulate negatively NFκB activity and prevent excessive inflammation ^42,43^. hnRNPL is traditionally an RNA processing factor that regulates alternative splicing by binding exonic splicing silencer elements resulting in exon exclusion from the mature mRNA^44,45^. The lncRNAs THRIL (TNF and hnRNPL related immunoregulatory lncRNA) and lincRNA-EPS both regulate inflammatory gene signaling via hnRNPL binding^46,47^. THRIL exerts its effects on TNF-α expression by binding to hnRNPL. LincRNA-EPS associates with chromatin in resting macrophages, repressing immune gene expression. Upon LPS stimulation, lincRNA-EPS in downregulated and hnRNPL is released from chromatin allowing for a more permissive chromatin state and immune genes transcription. Because of these novel inflammatory roles for the lncRNAs and hnRNPL, we thought to examine the interaction between it and PACERR.

**Figure 4.**
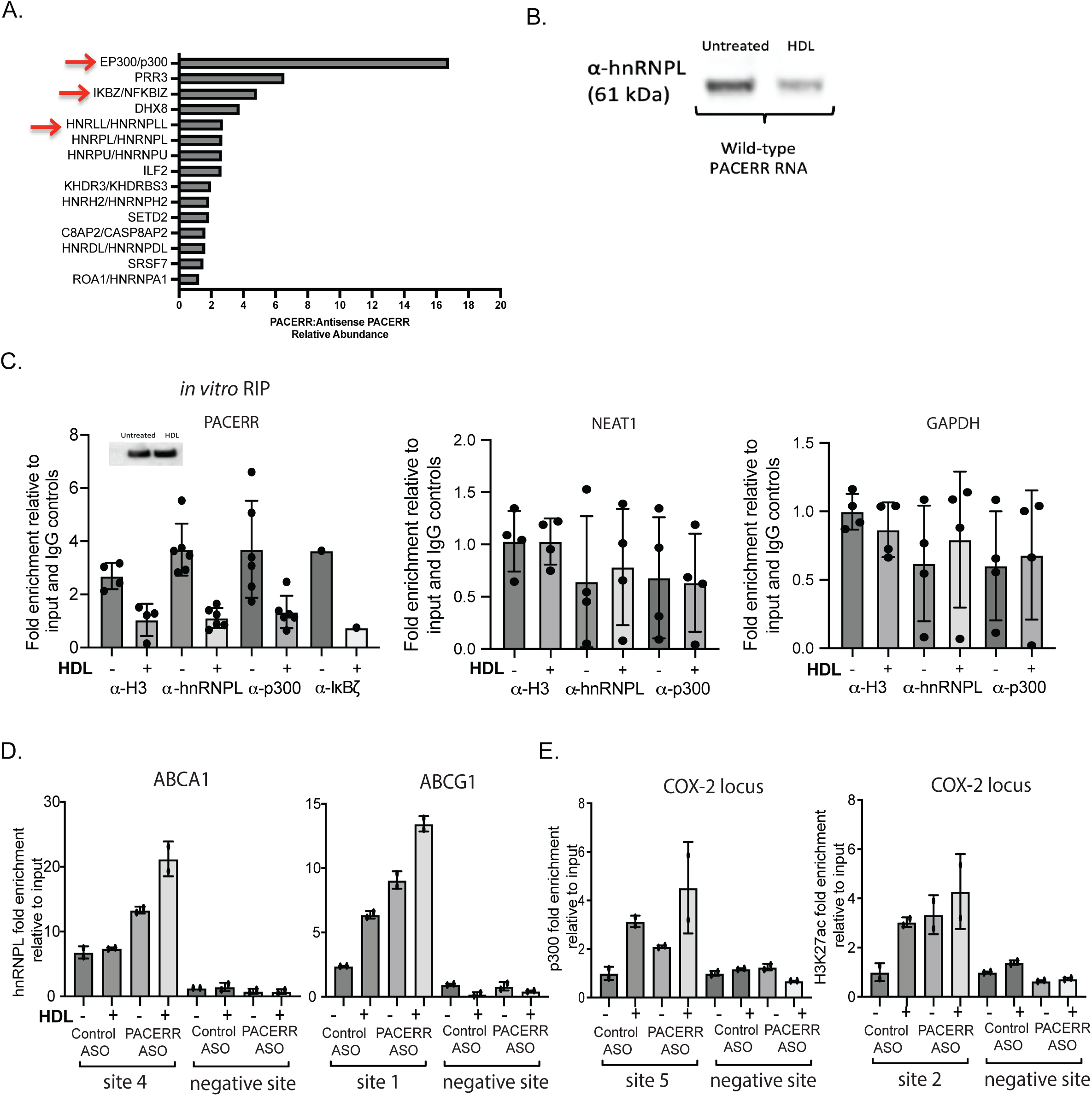
PACERR RNA interacts with p300, IκBζ and hnRNPL proteins basally and these interactions are attenuated upon HDL treatment. (A) PACERR cloned into pGEM-7Z+ for in vitro-transcription and biotin labeling. The biotinylated RNA was then incubated with nuclear cellular extracts from PMA differentiated human THP-1 cells. Fold change of abundance for proteins interacting with PACERR RNA versus PACERR antisense RNA are shown. (B) Western blot for hnRNPL is shown in unstimulated THP-1 macrophages or treated with 50µg/ml HDL for 24 hours. (C) THP-1 macrophages protein lysates either unstimulated or treated with 50µg/ml HDL for 24 hours were incubated with antibodies for H3, hnRNPL, p300 or IκBζ and qPCR for bound RNA was performed. NEAT1 and GAPDH were used as control for PACERR specificity. (n=6). (D) Chromatin immunoprecipitation for hnRNPL binding to ABCA1 and ABCG1 gene loci. (E) Enrichment for H3K27ac and p300 binding to the COX-2 gene locus. Fold enrichment relative to input is shown. Primers binding to region outside of the gene loci for each were used for negative controls. (n=2).

We confirmed hnRNPL association with PACERR using western blot. hnRNPL binds basally and upon HDL treatment, this interaction dissipates (Figure 4B). We confirmed protein-binding partners using RNA-immunoprecipitation (RIP) where untreated and HDL treated THP-1 macrophage protein lysates are enriched using antibodies for a specific protein and quantitative PCR (qPCR) is then used to measure RNAs that have bound to the proteins. We first measured enrichment of the H3 histone protein for PACERR to determine if PACERR is localized to chromatin, and then used antibodies for hnRNPL, p300 and IkBζ. We found in basal interactions for the proteins and the interactions decreased following HDL treatment (Figure 4C). NEAT1 lncRNA and GAPDH were used as negative controls.

Because of the effect of PACERR on cholesterol efflux and ABCA1 and ABCG1 expression, we then checked their genomic sequences for hnRNPL motifs as well as chromatin immunoprecipitation experiments (ChIP) experiments for hnRNPL enrichment in their loci. We found hnRNPL binding in intronic regions for both genes. We designed primers targeting these regions and then used the hnRNPL antibody to enrich chromatin for regions bound to the protein. We found hnRNPL bound to ABCA1 and ABCG1 in cells where PACERR was silenced with the ASO signifying PACERR acting as brake to negatively regulate the cholesterol transporters (Figure 4D).

Previous studies have shown p300 binding to the COX-2 promoter and affecting the activity of CREB in inflammatory settings and in the presence of HDL^38,48^. We wanted to see if there was enrichment of p300 and subsequent H3K27ac deposition at the COX-2 promoter following HDL treatment. We found p300 enrichment at the COX-2 promoter in response to HDL which was augmented when PACERR was silenced (Figure 4E). In agreement with the p300 increase, we also observed increased H3K27ac at the promoter signifying more accessible chromatin which leads to increased transcription of COX-2.

### The hnRNPL motif in PACERR RNA is required for ABCA1 and ABCG1 expression and cholesterol efflux

To further investigate the how PACERR is mediating its effects on hnRNPL binding to the genomic loci of ABCA1 and ABCG1, we examined the PACERR sequence for hnRNPL motifs. A strong binding site was found in the 825bp PACERR RNA sequence (Figure 5A). We confirmed the dependency on this site for the interaction we found in untreated versus HDL treated THP-1 macrophages and deleted the site from the PACERR sequence and found the interaction between PACERR and hnRNPL was abrogated (Figure 5B). We generated PACERR overexpressing THP-1 macrophage cell lines and deleted the hnRNPL binding site from the PACERR sequence (ΔhnRNPL). We found that when PACERR was overexpressed, hnRNPL binding to ABCA1 and ABCG1 was abrogated compared to empty vector (EV) control cells but in cells overexpressing PACERR with the hnRNPL site deleted, there was further enrichment of hnRNPL binding to the gene loci (Figure 5C).

**Figure 5.**
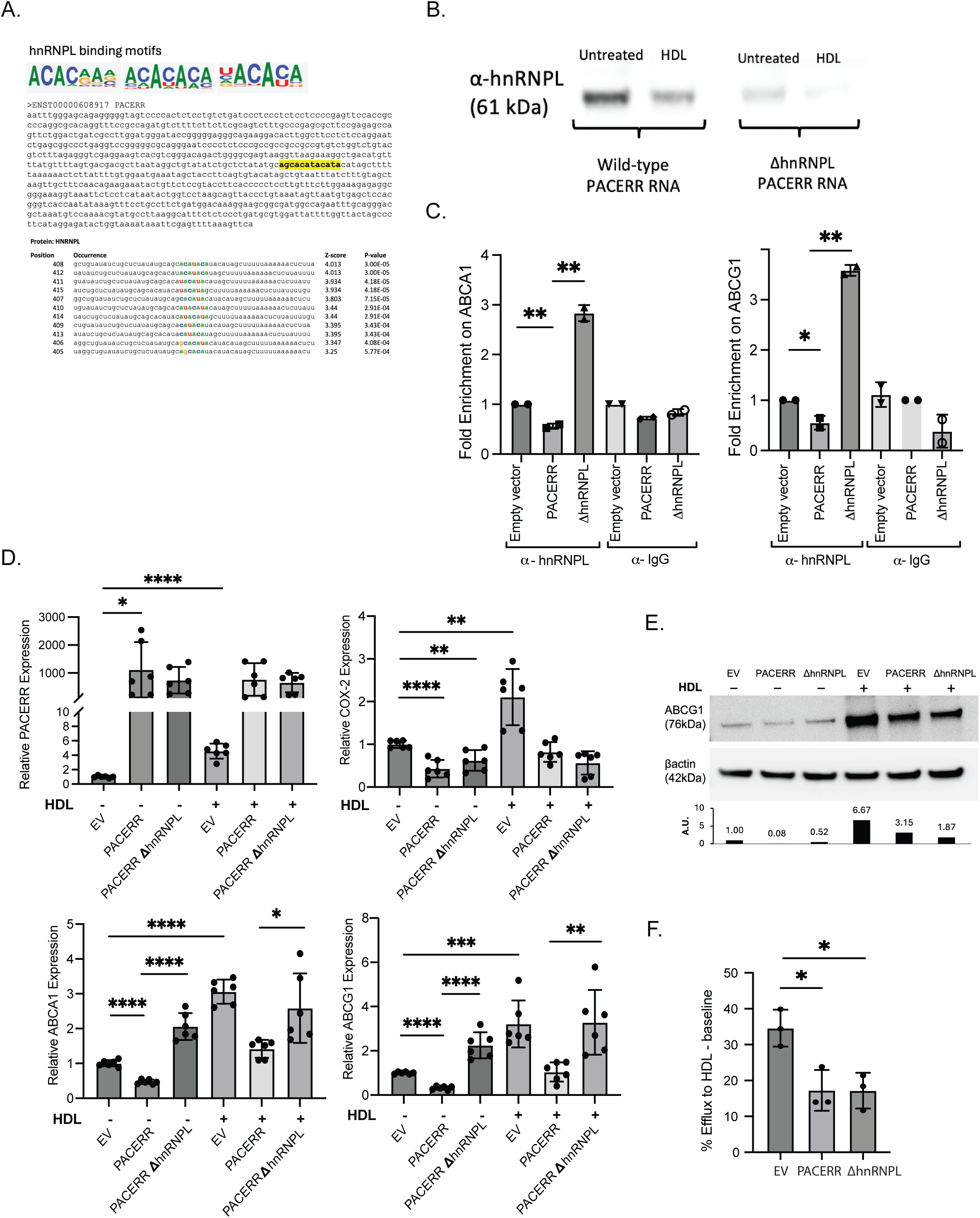
The hnRNPL motif is required for the PACERR mediated regulation of ABCA1 and ABCG1 and cholesterol efflux to HDL. (A) RBPMap identified an hnRNPL binding motif in the PACERR sequence, Z-scores for the strength of the motif are shown. (B) The hnRNPL motif was mutated in the PACERR pGEM-7Z+ plasmid for in vitro transcription. THP-1 macrophages were either unstimulated or treated with 50µg/ml HDL for 24 hours and protein lysates were harvested and incubated with wild-type PACERR RNA or ΔhnRNPL PACERR RNA. Western blot analysis is shown for hnRNPL levels. (C) Chromatin immunoprecipitation of THP-1 macrophages stably overexpressing PACERR or ΔhnRNPL PACERR using hnRNPL antibody and primers targeting the predicted hnRNPL binding site in ABCA1 and ABCG1. Fold enrichment relative to input control is shown. (n=2). (D) qPCR for PACERR, COX-2, ABCA1 and ABCG1 gene expression in THP-1 macrophages stably overexpressing PACERR or ΔhnRNPL PACERR and either unstimulated or treated with 50µg/ml HDL for 24 hours. (n=6) (E) ABCG1 protein levels shown by western blot in THP-1 macrophages stably overexpressing PACERR or ΔhnRNPL PACERR and either unstimulated or treated with 50µg/ml HDL for 24 hours. (F) Cholesterol efflux by THP-1 macrophages stably overexpressing PACERR or ΔhnRNPL PACERR following loading with ^3^H-cholesterol and 37.5µg/ml acLDL followed by HDL for efflux of cholesterol. Percent efflux is expressed as a percentage of total cell ^3^H-cholesterol content minus baseline efflux. (n=3). Values are mean ±SEM of at least three independent experiments. *p<0.05, **p<0.01, ***p<0.005, ****p<0.001 versus control untreated

Because of the known role for hnRNPL in alternative splicing, we then measured the expression of ABCA1 and ABCG1 in the PACERR overexpressing macrophages and ΔhnRNPL cells. We found PACERR expression increased over 1000-fold and caused decreased COX-2 expression, consistent with its sequestration of p300 from COX-2 leading to decreased transcription. ABCA1 and ABCG1 were also decreased, but we observed that in the ΔhnRNPL, their expression was restored to EV levels, consistent with the finding of increased hnRNPL binding to their loci (Figure 5D). However, when we measured ABCG1 protein level it remained decreased in both PACERR and ΔhnRNPL cells (Figure 5E) as well as cholesterol efflux levels, the functional output of ABCG1 (Figure 5F). This signifies that when PACERR no longer binds to hnRNPL, there is a defect in alternative splicing. mRNA expression levels are restored but the transcripts generated are not able to create a functioning protein required for proper cholesterol efflux to occur.

### PACERR interacts with cPLA2 to mediate LXR dependent ABCA1 and ABCG1

When HDL or LPS are applied to cells, we see an increase in the transcription of PACERR and a release of RNA binding proteins, p300, IkBζ and hnRNPL but we next questioned what this excess RNA could be doing in the cell. PLA2G4A is the neighboring gene of COX-2 and PACERR and is responsible for the release of arachidonic acid from cell membranes including nuclear membranes leading to the prostaglandin cascade, led by COX-2. HDL treatment of macrophages led to increased cPLA2 expression and was this was further increased in cells where PACERR was silenced with the ASO (Figure 6A). Consistent with this increase, we found cells released more arachidonic acid in the media. We found a similar result in ECs, where laminar flow induced cPLA2 expression and arachidonic acid release and HDL further increased this induction (Figure 6B). PACERR knockdown, further increased cPLA2 expression and arachidonic acid levels.

**Figure 6.**
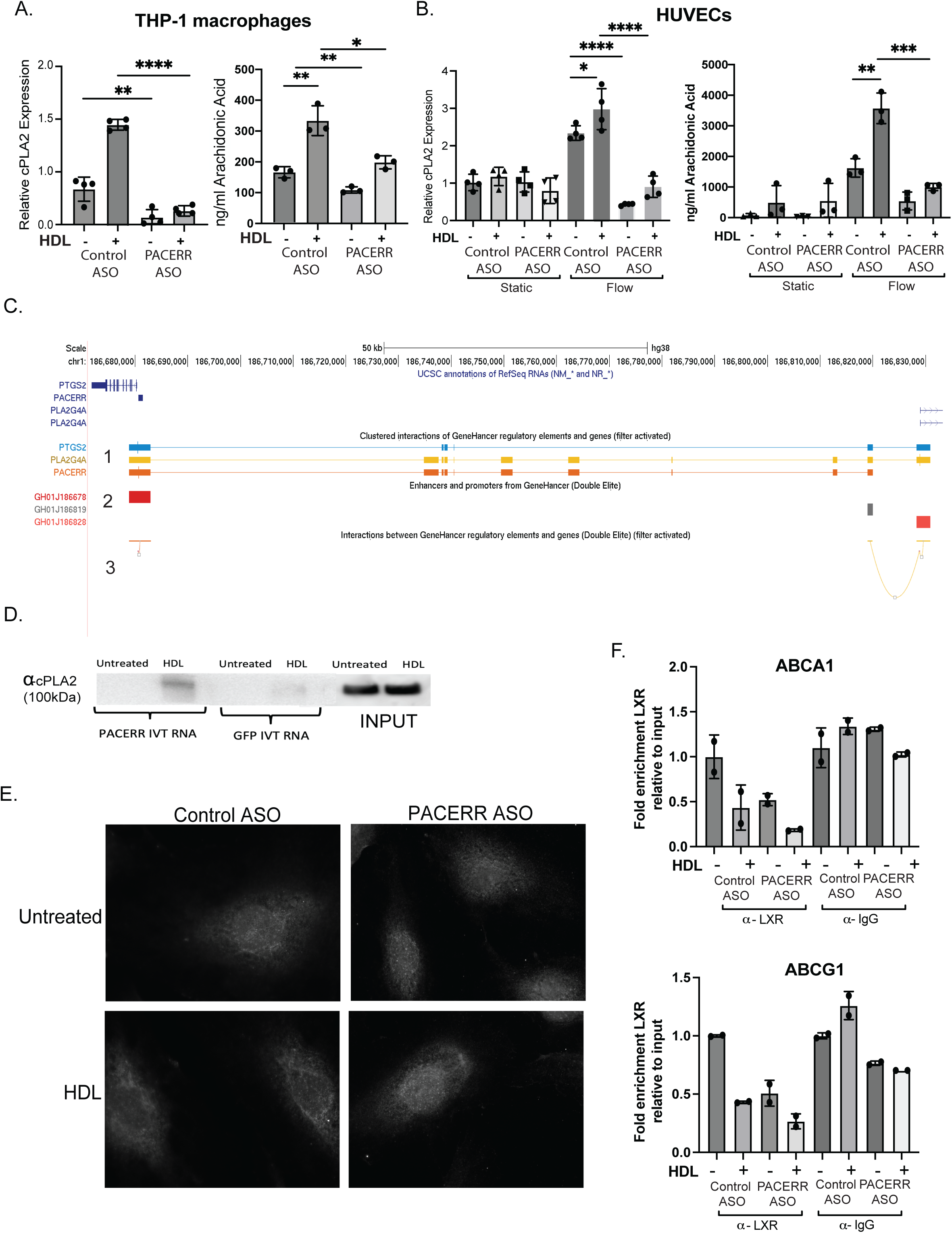
Upon HDL treatment, PACERR RNA interacts with cytosolic phospholipase A1 to mediate arachidonic acid release from the nuclear membrane. (A) qPCR for cPLA2 expression in THP-1 macrophages transfected with 50nm control or PACERR ASO and unstimulated or treated with 50µg/ml HDL for 24 hours. (n = 4). Arachidonic acid levels (ng/ml) in the media of the same cells. (B) qPCR for cPLA2 expression in HUVECs transfected with 50nm control or PACERR ASO and unstimulated or treated with 50µg/ml HDL for 24 hours under static or laminar flow (10 dyn/cm^2^) conditions. (n = 4). Arachidonic acid levels in the media of the same cells. (C) Genehancer track from Genome browser shows PACERR, PTGS2 and PLA2G4A interacting at a potential enhancer (gray box) that is mediating transcription of cPLA2 at its promoter region (red box). (D) Western blot showing cPLA2 protein interaction with PACERR *in vitro* transcribed RNA following treatment of THP-1 macrophages with 50µg/ml HDL for 24 hours. GFP in vitro transcribed RNA is used as a control. (E) THP-1 macrophages transfected with 50nm control ASO or PACERR ASO and stained for cPLA2 protein localization following HDL treatment. (F) Chromatin immunoprecipitation of THP-1 macrophages transfected with 50nm control or PACERR ASO followed by treatment with 50µg/ml HDL for 24 hours. LXR antibody and primers targeting the predicted LXR binding site in ABCA1 and ABCG1. Fold enrichment relative to input control is shown. (n=2).

To determine how PACERR could be mediating this effect on cPLA2, we next examined the PACERR/cPLA2 gene region (1) for chromatin/chromatin interactions using the Genehancer track on genome browser and found an enhancer region (2) where PACERR and cPLA2 chromatin are interacting and this enhancer further interacts with the cPLA2 promoter region showing its potential requirement for cPLA2 transcription (3 yellow line) (Figure 6C). We hypothesize that PACERR is required to activate cPLA2 transcription via regulation of the enhancer element and when it is silenced, it no longer can bring cPLA2 to its enhancer required for transcription.

We next found that upon HDL treatment, in vitro transcribed PACERR RNA and cPLA2 protein interact (Figure 6D) and observed using fluorescent microscopy that cPLA2 localizes to the nuclear membrane following HDL treatment and this is abrogated when PACERR is silenced (Figure 6E). This demonstrates the requirement of PACERR for escorting cPLA2 to the nuclear membrane where it is needed release HDL and flow induced arachidonic acid and subsequent PG formation. Previous studies have shown that arachidonic acid inhibits the binding of the LXR transcription factor to ABCA1 and ABCG1 ^4,49^. We found that HDL treatment of THP-1 macrophages led to reduced binding of LXR to the ABCA1 and ABCG1 promoters (Figure 6F). PACERR and cPLA2 are acting as an additional level of control over the LXR mediated induction of the cholesterol transporters. We have found that HDL regulates ABCA1 and ABCG1 at the level of alternative splicing by hnRNPL (Figure 7).

**Figure 7.**
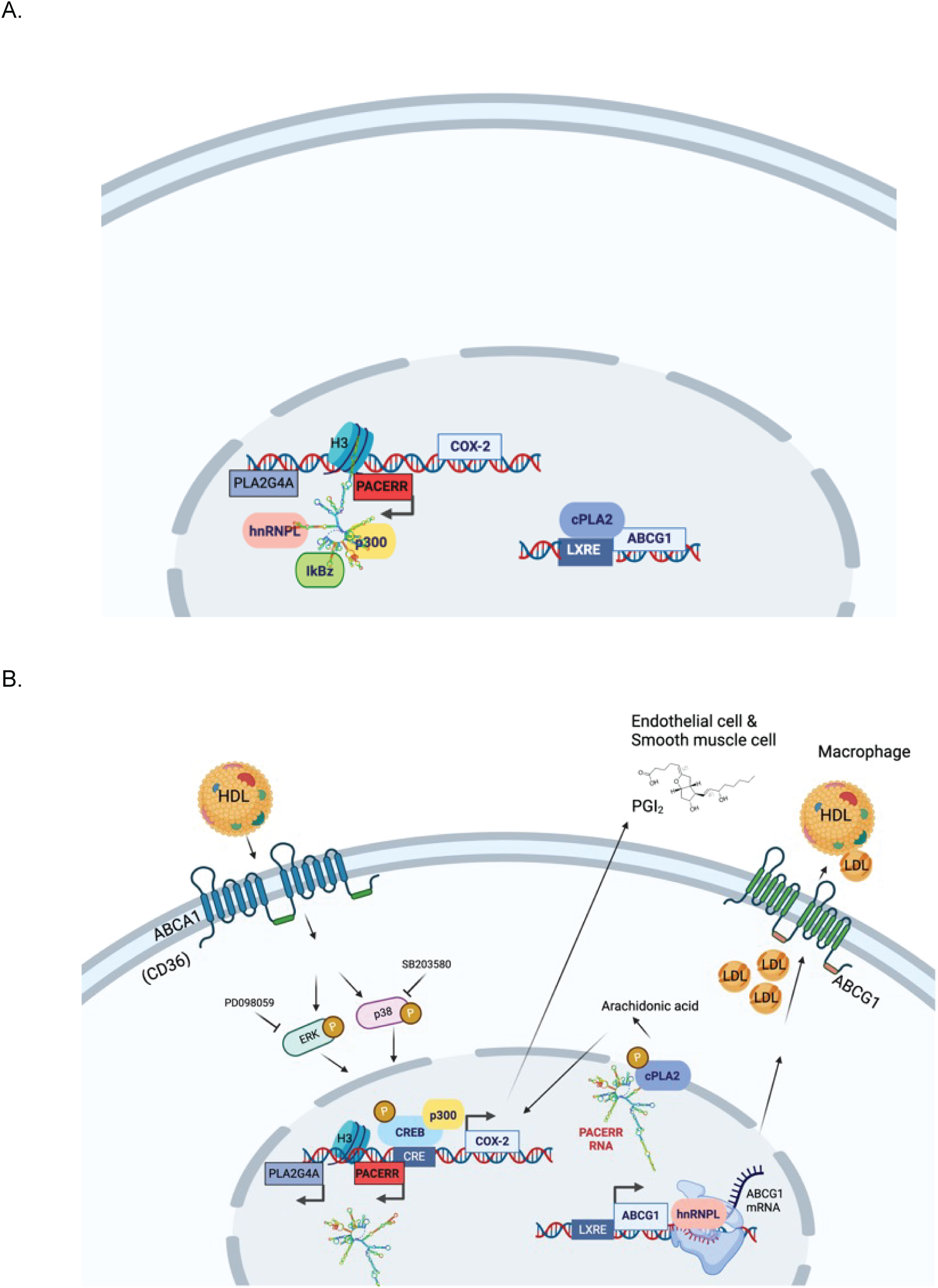
PACERR mediates PG synthesis and cholesterol efflux, linking two processes implicated in cardiovascular disease.

## Discussion

This study describes a novel mechanism for controlling the repertoire of PGs produced by cells in the vasculature. Following the observation that PACERR is primarily expressed in inflammatory macrophages and ECs in human carotid endarterectomies and its association with early, peripheral plaque formation, we focused on characterizing how PACERR controls COX-2 and the cholesterol transporters. Through simultaneous regulation of COX-2 transcription and alternative splicing of ABCA1 and ABCG1, the lncRNA PACERR acts to bridge PG synthesis with cholesterol efflux and thereby potentially influence atherogenesis.

PACERR, unlike the majority of annotated lncRNAs exhibits RNA expression in both mouse and human originating from the same genomic region. Primate and rodent COX-2 genes are highly conserved (∼89%) whereas the COX-2 promoter and PACERR region have diverged over time at a much more rapid pace. Only a 16-nucleotide NFκB site is maintained throughout primate and rodent lineages. In order to study the effects of human PACERR *in vivo* using models of acute inflammation and cholesterol transport, we generated a humanized mouse model where a 35kb mouse locus containing COX-2, PACERR and the region upstream were replaced with a 59kb syntenic human region.

We found that despite the overlap between the COX-2 promoter and PACERR, induction of PACERR by LPS or HDL is independent of COX-2 activity and unlike COX-2, which depends on NFκB, PACERR expression is solely dependent on the CREB transcription factor. We went on to identify protein binding partners for PACERR to elucidate its mechanism of action. We observed basal interactions between PACERR RNA, p300, IkBζ and hnRNPL and following cell stimulation with HDL or LPS, we saw a differential effect on the localization of the proteins. We were unable to detect an interaction between PACERR and p50 as previously described ^26^, but identified a basal interaction with IkBζ which preferentially associates with p50 versus p65 to regulate negatively NFκB activity and prevent excessive inflammation ^42,43^. LPS stimulated COX-2 expression, PG synthesis and systemic inflammation were all attenuated when PACERR was silenced in vivo because PACERR is no longer present to sequester IkBζ allowing it to bind to the NFκB site in COX-2, repressing its transcription. This is reminiscent of the regulation of Cox-2 expression by glucocorticoids, where the interaction between p65 and NFκB is also disrupted ^50^.

HDL has anti-atherogenic, anti-thrombotic and anti-inflammatory properties and is known to induce the expression of endothelial COX-2 dependent PGI_2_ production *in vitro*^51^. Platelet activation by ECs is a central aspect of atherogenesis and this can be inhibited by modulation of ApoA1 and HDL levels. The primary role for HDL is in facilitating RCT but it also functions to prevent the oxidation of LDL and subsequent accumulation of cholesterol^52^. However, HDL is a heterogeneous lipoprotein that consists of several functionally distinct subfractions. Small, dense HDL3 particles and large, light HDL2 particles differ in their ability to remove oxidized lipids from atherogenic LDL, preventing the accumulation of cholesterol and affecting the inflammatory status of the cell ^52-54^. Previous studies of ECs and vSMCs have shown a differential impact of HDL2 versus HDL3 on the expression of COX-2 and production of PGI_2_, potentially affording restraint on thrombogenesis, atherogenesis and hypertension^55^. ^38,56^. We found that subfractions of HDL also differentially influence *PACERR* expression in macrophages, with HDL3 inducing PACERR greater than HDL2. The subfractions also showed a differential effect on PG synthesis with HDL3 stimulation causing greater synthesis of PGE_2_ ^12,10^.

Cell exposure to exogenous HDL, led to the CREB-dependent transcriptional upregulation of PACERR. This was followed by the release of p300 from PACERR, its association with the CREB sites in the COX-2 promoter where its histone acetyltransferase activity caused increased H3K27ac marks and more accessible chromatin, all culminating in increased transcription of COX-2. Silencing PACERR in macrophages, vSMCs and ECs led to augmented COX-2 expression and PG production in response to HDL, while also increasing the expression of ABCA1 and ABCG1 and cholesterol efflux to HDL. We found that PACERR controls the levels of the cholesterol transporters through its interaction with the splicing factor, hnRNPL. It is released from PACERR RNA in response to HDL where it relocates to introns of ABCA1 and ABCG1 acting in the mediator complex and determining the alternative splicing fate of transcripts required for cholesterol efflux. The binding of hnRNPL to the ABCA1 and ABCG1 genes is augmented when PACERR is silenced. Illustrating the specificity for the interaction between hnRNPL and PACERR, we also found when the hnRNPL motif in the PACERR gene sequence is deleted, it no longer binds and is free to associate with the cholesterol transport genes, increasing their transcription and expression levels.

We observed an additional layer of regulation over the cholesterol efflux pathway through the association between PACERR and its neighboring gene, cPLA2/PLA2G4A, an enzyme responsible for the release of arachidonic acid from the nuclear membrane. Arachidonic acid inhibits the formation of the LXR/RXR heterodimer required for the transcription of ABCA1 and ABCG1 ^4,57^. Following treatment with HDL, cPLA2 translocates to the nuclear membrane, binds to PACERR, and releases arachidonic acid. This sequence was disrupted by silencing PACERR, whereupon LXR bound to the promoters of ABCA1 and ABCG1 increasing their expression. This study illustrates multiple layers of control over cholesterol metabolism, involving splicing of cholesterol transport transcripts as well as transcriptional regulation via LXR.

These findings advance our understanding of how a single lncRNA can mediate the cell specific response to HDL and LPS but raises future questions to further investigate how PACERR is acting as a fine tuner of the inflammatory response through its interactions with an assembly of protein binding partners to mediate pathways involved in acute and potentially chronic diseases. There was a basal interaction between PACERR, p300, IkBζ and hnRNPL and observed that cell stimulation causes active transcription of PACERR which will result in changes to its RNA secondary structure compared to that of basal PACERR RNA. We anticipate that his change in RNA structure allows for the differential release of p300, IkBζ and hnRNPL to relocate to the COX-2 promoter and the introns of ABCA1 and ABCG1. Our mechanistic studies demonstrate key players in mediating the response to LPS or HDL but, it will be of interest to determine if LPS or HDL alter the secondary structure of PACERR differently to control the fate of the protein binding partners.

The maintenance of lipid homeostasis is tightly regulated involving various signaling pathways where disruptions in a pathway such as from causal single nucleotide polymorphisms (SNPs) may result in hyperlipidemia and atherosclerosis. It will be of interest to interrogate the human genetics associated with PACERR expression. Because we found evidence for PACERR in ECs and inflammatory macrophages in human carotid plaques, implicating it in endothelial dysfunction and that treating acute inflammation with a PACERR ASO reduced inflammation, we anticipate that people with elevated PACERR expression may be at risk for NSAID induced cardiovascular complication and the development of atherosclerosis.

Previous studies have demonstrated the importance of COX-2 in mediating atherogenesis. Its expression is induced by HDL and LPS to differentially produce PGs involved in vascular inflammation and platelet function. By identifying a common transcriptional program where PACERR is induced by both LPS and HDL via CREB signaling, but with diverging functional outcomes, we have elucidated a cell specific response for maintaining cellular homeostasis. Here we’ve shown a new mechanism whereby the differential functionalities of HDL and LPS are linked through the augmentation or restraint of vascular inflammation. Through various lines of evidence, we have integrated PG formation and cholesterol metabolism through PACERR. Extracellular cholesterol uptake and efflux along with endogenous fatty acid biosynthesis pathways are responsible for regulating lipid levels and inflammation and a greater understanding of how these signaling pathways are controlled by lncRNAs may contribute to novel therapeutics for the prevention of cardiovascular disease and cardiotoxicity associated with NSAID use.

## Materials & Methods

### Ethics statement

The investigators faithfully adhered to the “Guide for the Care and Use of Laboratory Animals” by the Committee on Care of Laboratory Animal Resources Commission on Life Sciences, National Research Council. The animal facilities at the University of Pennsylvania are fully accredited by the American Association for Accreditation of Laboratory Animal Care (AAALAC). All studies were conducted under protocols approved by the University of Pennsylvania IACUC.

### Human studies

See Paloschi *et al* for processing and single cell RNA sequencing of human carotid endarterectomy samples ^28^.

### Mouse studies

Mice were kept under 12:12 light:dark conditions with environmental enrichment (nestlet, shepherd shacks) unless otherwise specified. All mice were housed with ad libitum access to standard chow diet and water. Studies were performed in accordance with the protocol approved by the University of Pennsylvania Institutional Animal Care and Use Committee. More details are given in the Supplementary Materials.

### Cell culture

HEK293T and THP-1 cell lines were obtained from American Type Tissue Collection, authenticated with standard American Type Tissue Collection methods (morphology check under a microscope and growth curve analysis) and tested monthly for mycoplasma contamination. HEK293T were maintained in high-glucose DMEM (Corning) supplemented with 10% fetal bovine serum (FBS, Life Technologies) and 1% penicillin–streptomycin (Life Technologies). THP-1 cells were maintained in RPMI 1640 (Gibco, 11875093) with 10% FBS, 1% penicillin–streptomycin (100×, Gibco, 15140122), 10 mM Hepes, 1 mM Sodium pyruvate, and 0.05 mM ß-mercaptoethanol according to ATTC and were differentiated into macrophages in the presence of 10 nM phorbol-12-myristate acetate (PMA, Sigma) for 48–72 h. Vascular aortic smooth muscle cells were obtained from Lonza from 3 different male donors (cat # CC-2571). vSMCs were grown in SmGM-2 Smooth Muscle Cell Growth Medium-2 BulletKit (cat # CC-3182). Human umbilical vein endothelial cells (HUVECs) were obtained from Lonza (cat # CC-2519) and grown in EGMTM Endothelial Cell Growth Medium BulletKit (cat # CC-3124).

### Culture of primary cells

Primary peripheral blood mononuclear cells (PBMCs) were isolated from blood collected in a BD vacutainer CPT (REF 362753). Whole blood was centrifuged through a Ficoll-Paque PLUS (GE Healthcare) density gradient centrifugation, and PBMCs were collected from the buffy coat. Macrophages were expanded and cultivated in the presence of 50 ng/mL recombinant human M-CSF (R&D, 216-MC-025) to generate monocyte derived macrophages (MDMs). Cells were grown in RPMI-1640, 1% penicillin/streptomycin, 10% FBS, 25mM Hepes, 1X nonessential amino acids (100×, Life Sciences, SH3023801), 1X GlutaMax (100×, Gibco, 35050-061), 1mM Na-pyruvate (Gibco, 11360-070), 10µg ciprofloxacin (Acros, AC456880050). Mouse bone marrow–derived macrophages (BMDMs) were isolated from femurs and tibias as described previously^58^. Macrophages were differentiated in DMEM supplemented with 10% FBS, 1% penicillin/streptomycin and 15% L929 media. Cells were incubated at 37°C in 5% CO2.

### Cellular response to lipoproteins and inflammatory ligands

Apolipoprotein A1 (ApoA1) was purchased from molecular cloning laboratories (cat # APO-100) and used at 50µg/ml. High density lipoproteins (HDL, HDL2 and HDL3) were purchased from Lee biosolutions (cat # 361-10-0.1) or academy biomedical solutions (80P-HD-101, 80P-HD2-101, 80P-HD3-101) and used at 50µg/ml, acetylated LDL (acLDL) was purchased from Kalen Biomedical, LLC (770201-6) and used at 37.5µg/ml. Oxidized LDL was purchased from Lee Biosolutions (360-31-1) and used at X. Ultrapure Lipopolysaccharides from *Escherichia coli* O111:B4 (LPS) was used at 100ng/ml. Cells were incubated for 8 or 24 hours. Cells were pretreated with celecoxib (Carbosyth FC1130) or Bay 11-7082 (Sigma B5556).

### RNA isolation, cell fractionation and real-time PCR

Total RNA was isolated using TRIzol reagent (Invitrogen) and Direct-zol RNA MicroPrep columns (Zymo Research). Tissues were lysed with steel beads using the Qiagen tissue lyser in the presence of Trizol. For cell fractionation experiments RNA was isolated from separate cytoplasmic and nuclear fractions using the RNA Subcellular Isolation Kit (Active Motif 25501). Upon isolation, RNA was reverse transcribed using Applied biosystems cDNA Synthesis kit using multiscribe reverse transcriptase (Life tech 4311235) and quantitative PCR analysis was conducted using Applied biosystems FAST SYBR green (4385612) according to the manufacturer’s instructions and quantified on the ViiA 7 Real-Time PCR System (Applied Biosystems). Fold change in mRNA expression was calculated using the comparative cycle method (2−ΔΔCt) normalized to the housekeeping gene HPRT. A list of primers used in this study can be found in supplement.

### Gain- and loss-of-function studies

Transient knockdown of CREB was done by designing shRNAs targeting a region to all CREB gene variants or a non-targeting control shRNA and cloned into the pLKO vector (Addgene, 8453) and transfected into HEK293T cells with packaging vectors psPAX2 (Addgene, 12260) and pMD2.G (Addgene, 12259). After 48 h, the culture medium was isolated and used to transduce THP-1 cells. Cells stably expressing the shRNAs were selected by using 5 μg ml^−1^ puromycin (Thermo Fisher Scientific). To create PACERR-overexpressing cell lines, PACERR gene block from IDT was cloned into the pMSCV-PIG vector (Addgene, 21654) and transfected into HEK293T cells with packaging vectors pCMV-VSV-G (Addgene, 8454) and pCMV-Gag-Pol (Cell Biolabs, RV-111). After 48 h, the culture medium was isolated and used to transduce THP-1 cells were selected with puromycin (5 μg ml^−1^, Thermo Fisher Scientific).

For knockdown of PACERR in PMA-differentiated THP-1 cells, 50 nM locked nucleic acid GapmeRs (IDT) targeting PACERR or Negative Control GapmerRs (CTRL) were transfected using Lipofectamine RNAiMax (Life Technologies) as described^24^. Knockdown of ABCA1 and ABCG1 in THP-1 cells was acquired by transfecting 50 nM Dharmacon ON-TARGET plus siRNAs directed against ABCA1 (J-004128-06-0002) or ABCG1 (J-008615-05-0002) or non-targeting control (D-001810-01-05) using Lipofectamine RNAiMax. After the 24 hours of incubation, cells were treated with HDL and then lysed and RNA was extracted.

To mutate the predicted binding sites for hnRNPL, we deleted the motif with a QuikChange XL kit (Stratagene), by using the primers indicated in Supplementary Table. PACERR^MUT^-expressing cell lines were generated as described above. All constructs were confirmed by sequencing.

### Lipid nanoparticle preparation and characterization

LNPs used in this study were similar in composition to those described previously^59^, an ionizable cationic lipid (proprietary to Acuitas)/phosphatidylcholine/cholesterol/PEG-lipid (50:10:38.5:1.5 mol/mol) SM-102 was used. The nitrogen to phosphate ratio (N/P ratio) was 5 and they were encapsulated at an RNA to total lipid ratio of ∼0.05 (wt/wt). Formulation was prepared using microfluidic mixing of lipids (in ethanol) and ASO (in sodium acetate), and buffer exchanged against DPBS buffer supplemented with 8% w/v of sucrose. They had a diameter of ∼80 nm as measured by dynamic light scattering using a Zetasizer Nano ZS (Malvern Instruments Ltd., Malvern, UK) instrument. mRNA-LNP formulations were stored at −80 °C at a concentration of mRNA of ∼1 μg/μL. LNPs were conjugated with mAb specific for PECAM-1. Targeting antibodies or control isotype-matched IgG were conjugated to LNP particles *via* SATA–maleimide conjugation chemistry using the previously established method ^59^.

### Application of shear stress

HUVECs were seeded on gelatin-coated μ-slides 0.6 Luer (Ibidi 50-305-756) for 3 hours and transfected with 10nm LNP-ASOs. Flowing medium was applied by using the ibidi pump system to generate unidirectional laminar flow (10 dyn/cm^2^), oscillating flow (0.5 + 4dyn/cm2), or static control for 24 hours in an incubator. The system was maintained at 37 °C and ventilated with 95% humidified air containing 5% CO2.

### Preparation of cells for chromatin immunoprecipitation (ChIP)

ChIP-immunoprecipitation was performed using SimpleChIP Plus Sonication Chromatin IP Kit (#56383, Cell Signaling Technology) following manufacturer’s instructions. Briefly, cells were cross-linked with 37% formaldehyde for 10 min at room temperature. Isolated chromatin was fragmented using the Diagenode Bioruptor (ON 30 sec, OFF 30 sec) for 30 mins to generate 200 – 500 bp chromatin fragments. In each experiment, chromatin was first processed by agarose gel electrophoresis to confirm DNA shearing to 200 - 500 bp fragments, and the DNA concentration was measured by NanoDrop 2000. Equal quantities of sheared chromatin (10 μg per immunoprecipitation) were diluted 1:5 in sonication buffer to the final volume of 1 mL, and immunoprecipitated overnight with 1 μg antibody targeting human hnRNPL (novus NB120-6106), P300 (Abcam ab275378), Histone H3K27Ac (Active Motif, 39685), IkBζ (Cell Signaling 9244S), cPLA2 (Cell Signaling 2832S) and LXR-α and LXR-β (Active motif 61175 and 61177). An equal amount of chromatin from each sample was immunoprecipitated with normal rabbit IgG (#2729P) at 4°C overnight. Chromatin complexes were captured using 20 μL Dynabeads protein G (Invitrogen) at 4°C for 1h.

### Western blots

Protein was extracted in RIPA buffer (Cell Signaling) with protease and phosphatase inhibitors (Roche) and subsequently normalized with a Pierce BCA Protein Assay Kit (Thermo Fisher Scientific). Samples (30 μg per well) were electrophoresed on 4–20% TGX-gradient gels (Bio-Rad Laboratories) and transferred to nitrocellulose membranes at 125 V for 2 h. Membranes were incubated overnight with the specified antibodies directed against COX-2 (Cayman 160106) ABCA1 (Novus Biologicals, NB400-105), ABCG1 (Novus NB400-132) and ß-actin (Sigma A5441). Proteins were visualized by using appropriate secondary antibodies and scanned with an Odyssey Imaging System (Li-Cor Biosciences). Quantification was performed in Image Studio software (Li-Cor Biosciences).

### Cholesterol efflux assay

For cholesterol efflux assays, THP-1 cells stably expressing PACERR or transfected with PACERR ASOs were seeded in 24-well plates at a density of 1 × 10^6^ cells, and THP-1 cells were treated with PMA as described above. Cells were labeled with 0.5 μCi ml^−1^ of ^3^H-cholesterol (PerkinElmer) and 37.5 μg ml^−1^ acetylated LDL (Kalen Biomedical, LLC 770201-6) for 24 h, washed twice with PBS and then incubated with 2 mg ml^−1^ fatty-acid-free BSA (Sigma-Aldrich) in medium for 24 h. To induce efflux, we added 50 μg ml^−1^ human HDL (Lee biosolutions 361-10-0.1) in fatty-acid-free medium, and supernatants were collected after 24 h. HDL-dependent efflux was expressed as a percentage of total cell ^3^H-cholesterol content.

### Mass spectrometry screen for RNA Binding Proteins

NE-PER Nuclear and Cytoplasmic Extraction Reagents (thermos 78833) was used to isolate nuclear proteins from THP-1 macrophages. We used PACERR cloned into pGEM-7Z+ (Promega) as the template to in vitro-transcribe (IVT) using T7 RNA polymerase, and biotin-16-UTP was incorporated at approximately every 20th to 25th nucleotide of the transcript.

Pierce™ Magnetic RNA-Protein Pull-Down Kit (Thermo 20164) was used to identify proteins bound to PACERR IVT RNA. Biotinylated RNA was incubated with nuclear cellular extracts from THP-1 macrophages unstimulated or treated with 50µg/ml HDL. Streptavidin magnetic beads were used to pull-down biotinylated PACERR and proteins bound to PACERR were identified using liquid chromatography-mass spectrometry (LC-MS) as previously described^60^. We used an antisense PACERR RNA sequence and GFP sequence as controls for specificity of the in vitro transcribed product. GFP is approximately the same size as the human PACERR transcript, 715bp versus 825bp.

### Measurement of HDL Paraoxonase activity and F2-Isoprostane levels

Paraoxonase or PON1 activity was measured by hydrolysis of paraoxon. 10 µL of the HDL sub-fractions were added to 95 µL buffer solution (100 mM Tris-HCl, 2 mM CaCl^2^, pH 8.0) and then 95 µl of 5.5 mM paraoxon-ethyl (Sigma-Aldrich, 36186) (190ul +10ul = 200ul reaction). Absorbance was recorded at 412 nm and 25°C over a period of 12 min (at 20-s intervals). Formation of *p*-nitrophenol /4-nitrophenol was measured. Blanks contained substrate without enzyme and were used to correct for the spontaneous hydrolysis of the substrate. Enzyme activity was calculated with a molar extinction coefficient of 18,290 M^−1^ cm^−1^. One unit of paraoxonase activity produced 1 nmol of p-nitrophenol per minute. 1 nmol of paraoxon hydrolyzed per ml per minute.

F2-isoprostanes present on HDL subfraction molecules were measured using LC-MS. SPIKE protein concentration was 0.1ng/uL for d4-iPF2a-III and d11-8,12-iso-iPF2a-VI. 50 uL SPIKE (5ng d4-iPF2a-III and 5ng d11-iPF2a-VI) was added to 100 uL sample. Then 10 ul 5% D,L-dithiothreitol solution was added to samples along with 100 μl of 15% KOH. The mixture was incubated at 40°C for 1 h. The solution was then acidified to pH 3 with 6N HCl. (∼50 uL). 700 uL water was added.

### Mass spectrometric snalysis of cell culture media prostanoids

Prostanoids in cell culture media were measured, as described by Meng et al^61^.

### Mass spectrometric analysis of urinary prostaglandin metabolites

Urinary prostanoid metabolites were measured by liquid chromatography/mass spectrometry/mass spectrometry as described^62^. Such measurements provide a noninvasive, time-integrated measurement of systemic prostanoid biosynthesis^63^.

### RNA Immunoprecipitation

Human histone H3 (Cell signaling), hnRNPL, p300, IkBζ and cPLA2 (antibodies used as above in ChIP) were immunoprecipitated from PMA-differentiated THP-1 macrophages. All immunoprecipitations were done using the MagnaRIP RNA-Binding Protein Immunoprecipitation Kit (EMD Millipore) according to the manufacturers’ instructions. Briefly, an antibody targeting protein of interest, or an isotype matched control IgG antibody were bound to magnetic beads and incubated with lysed cells at 4°C for 24h. Beads were isolated and cleaved from the bound proteins by proteinase K, and coprecipitated RNA was purified. qPCR analysis of total RNA was performed to detect enrichment of CHROMR variants and control genes in the protein-of-interest precipitated fraction was determined as percentage of 1% input control.

### Fluorescent microscopy

cPLA2 (Novus AF6659) was visualized using fluorescent donkey anti-goat 546 secondary antibodies (fisher A11056) and DAPI (Sigma D9542) was used to visualize nuclear DNA.

### Statistics

All animals compared were the stated age. Post hoc analysis was performed by pairwise *t* tests. All significant tests were further validated by the nonparametric Mann-Whitney *U* and Wilcoxon tests to make sure that they were not biased by parametric assumption. A significance threshold of 0.05 was used for all tests after correcting for multiple tests. Sample sizes were based on power analysis of the test measurement and the desire to detect a minimal 10% difference in the variables assessed with α=0.05 and the power (1−β)=0.8.

## Notes

### Competing Interest Statement

The authors have declared no competing interest.

## References

1. Crofford LJ, Wilder RL, Ristimaki AP, Sano H, Remmers EF, Epps HR, Hla T. Cyclooxygenase-1 and -2 expression in rheumatoid synovial tissues. Effects of interleukin-1 beta, phorbol ester, and corticosteroids. The Journal of clinical investigation. 1994;93:1095-1101. doi: 10.1172/JCI117060

2. Schonbeck U, Sukhova GK, Graber P, Coulter S, Libby P. Augmented expression of cyclooxygenase-2 in human atherosclerotic lesions. The American journal of pathology. 1999;155:1281–1291. doi: 10.1016/S0002-9440(10)65230-3

3. Narasimha A, Watanabe J, Lin JA, Hama S, Langenbach R, Navab M, Fogelman AM, Reddy ST. A novel anti-atherogenic role for COX-2--potential mechanism for the cardiovascular side effects of COX-2 inhibitors. Prostaglandins & other lipid mediators. 2007;84:24–33. doi: 10.1016/j.prostaglandins.2007.03.004

4. Zhou L, Choi HY, Li WP, Xu F, Herz J. LRP1 controls cPLA2 phosphorylation, ABCA1 expression and cellular cholesterol export. PloS one. 2009;4:e6853. doi: 10.1371/journal.pone.0006853

5. Ricciotti E, FitzGerald GA. Prostaglandins and inflammation. Arteriosclerosis, thrombosis, and vascular biology. 2011;31:986-1000. doi: 10.1161/ATVBAHA.110.207449

6. Massy ZA, Swan SK. Cyclooxygenase-2 and atherosclerosis: friend or foe? Nephrology, dialysis, transplantation : official publication of the European Dialysis and Transplant Association - European Renal Association. 2001;16:2286–2289.

7. Zhou W, Hashimoto K, Goleniewska K, O’Neal JF, Ji S, Blackwell TS, Fitzgerald GA, Egan KM, Geraci MW, Peebles RS, Jr. Prostaglandin I2 analogs inhibit proinflammatory cytokine production and T cell stimulatory function of dendritic cells. Journal of immunology. 2007;178:702–710.

8. FitzGerald GA, Smith B, Pedersen AK, Brash AR. Increased prostacyclin biosynthesis in patients with severe atherosclerosis and platelet activation. The New England journal of medicine. 1984;310:1065–1068. doi: 10.1056/NEJM198404263101701

9. Egan KM, Lawson JA, Fries S, Koller B, Rader DJ, Smyth EM, Fitzgerald GA. COX-2-derived prostacyclin confers atheroprotection on female mice. Science. 2004;306:1954–1957. doi: 10.1126/science.1103333

10. Hui Y, Ricciotti E, Crichton I, Yu Z, Wang D, Stubbe J, Wang M, Pure E, FitzGerald GA. Targeted deletions of cyclooxygenase-2 and atherogenesis in mice. Circulation. 2010;121:2654–2660. doi: 10.1161/CIRCULATIONAHA.109.910687

11. Yu Z, Crichton I, Tang SY, Hui Y, Ricciotti E, Levin MD, Lawson JA, Pure E, FitzGerald GA. Disruption of the 5-lipoxygenase pathway attenuates atherogenesis consequent to COX-2 deletion in mice. Proceedings of the National Academy of Sciences of the United States of America. 2012;109:6727–6732. doi: 10.1073/pnas.1115313109

12. Tang SY, Monslow J, Todd L, Lawson J, Pure E, FitzGerald GA. Cyclooxygenase-2 in endothelial and vascular smooth muscle cells restrains atherogenesis in hyperlipidemic mice. Circulation. 2014;129:1761–1769. doi: 10.1161/CIRCULATIONAHA.113.007913

13. Frikke-Schmidt R, Nordestgaard BG, Stene MC, Sethi AA, Remaley AT, Schnohr P, Grande P, Tybjaerg-Hansen A. Association of loss-of-function mutations in the ABCA1 gene with high-density lipoprotein cholesterol levels and risk of ischemic heart disease. Jama. 2008;299:2524–2532. doi: 10.1001/jama.299.21.2524

14. Voight BF, Peloso GM, Orho-Melander M, Frikke-Schmidt R, Barbalic M, Jensen MK, Hindy G, Holm H, Ding EL, Johnson T, et al. Plasma HDL cholesterol and risk of myocardial infarction: a mendelian randomisation study. Lancet. 2012;380:572–580. doi: 10.1016/S0140-6736(12)60312-2

15. Eliopoulos AG, Dumitru CD, Wang CC, Cho J, Tsichlis PN. Induction of COX-2 by LPS in macrophages is regulated by Tpl2-dependent CREB activation signals. The EMBO journal. 2002;21:4831–4840.

16. Wen AY, Sakamoto KM, Miller LS. The role of the transcription factor CREB in immune function. Journal of immunology. 2010;185:6413–6419. doi: 10.4049/jimmunol.1001829

17. Tong X, Lv P, Mathew AV, Liu D, Niu C, Wang Y, Ji L, Li J, Fu Z, Pan B, et al. The compensatory enrichment of sphingosine -1- phosphate harbored on glycated high-density lipoprotein restores endothelial protective function in type 2 diabetes mellitus. Cardiovascular diabetology. 2014;13:82. doi: 10.1186/1475-2840-13-82

18. Xiong SL, Liu X, Yi GH. High-density lipoprotein induces cyclooxygenase-2 expression and prostaglandin I-2 release in endothelial cells through sphingosine kinase-2. Molecular and cellular biochemistry. 2014;389:197–207. doi: 10.1007/s11010-013-1941-y

19. Vinals M, Martinez-Gonzalez J, Badimon JJ, Badimon L. HDL-induced prostacyclin release in smooth muscle cells is dependent on cyclooxygenase-2 (Cox-2). Arteriosclerosis, thrombosis, and vascular biology. 1997;17:3481–3488.

20. Dinger ME, Amaral PP, Mercer TR, Pang KC, Bruce SJ, Gardiner BB, Askarian-Amiri ME, Ru K, Solda G, Simons C, et al. Long noncoding RNAs in mouse embryonic stem cell pluripotency and differentiation. Genome research. 2008;18:1433–1445. doi: 10.1101/gr.078378.108

21. Jia H, Osak M, Bogu GK, Stanton LW, Johnson R, Lipovich L. Genome-wide computational identification and manual annotation of human long noncoding RNA genes. Rna. 2010;16:1478–1487. doi: 10.1261/rna.1951310

22. Ponjavic J, Oliver PL, Lunter G, Ponting CP. Genomic and transcriptional co-localization of protein-coding and long non-coding RNA pairs in the developing brain. PLoS genetics. 2009;5:e1000617. doi: 10.1371/journal.pgen.1000617

23. Ponjavic J, Ponting CP, Lunter G. Functionality or transcriptional noise? Evidence for selection within long noncoding RNAs. Genome research. 2007;17:556–565. doi: 10.1101/gr.6036807

24. Hennessy EJ, van Solingen C, Scacalossi KR, Ouimet M, Afonso MS, Prins J, Koelwyn GJ, Sharma M, Ramkhelawon B, Carpenter S, et al. The long noncoding RNA CHROME regulates cholesterol homeostasis in primate. Nature metabolism. 2019;1:98–110. doi: 10.1038/s42255-018-0004-9

25. van Solingen C, Cyr Y, Scacalossi KR, de Vries M, Barrett TJ, de Jong A, Gourvest M, Zhang T, Peled D, Kher R, et al. Long noncoding RNA CHROMR regulates antiviral immunity in humans. Proceedings of the National Academy of Sciences of the United States of America. 2022;119:e2210321119. doi: 10.1073/pnas.2210321119

26. Krawczyk M, Emerson BM. p50-associated COX-2 extragenic RNA (PACER) activates COX-2 gene expression by occluding repressive NF-kappaB complexes. eLife. 2014;3:e01776. doi: 10.7554/eLife.01776

27. Pauli J, Daniel Garger, Fatemeh Peymani, Justus Wettich, Nadja Sachs, Johannes Wirth, Katja Steiger, Hanrui Zhang, Ira Tabas, Alan Tall, Mingyao Li, Muredach P. Reilly, Daniela Branzan, Holger Prokisch, Michael P. Menden, Lars Maegdefessel. Single cell spatial transcriptomics integration deciphers the morphological heterogeneity of atherosclerotic carotid arteries. bioRxiv : the preprint server for biology. 2025.

28. Paloschi V, Pauli J, Winski G, Wu Z, Li Z, Botti L, Meucci S, Conti P, Rogowitz F, Glukha N, et al. Utilization of an Artery-on-a-Chip to Unravel Novel Regulators and Therapeutic Targets in Vascular Diseases. Advanced healthcare materials. 2023:e2302907. doi: 10.1002/adhm.202302907

29. Ramsay RG, Ciznadija D, Vanevski M, Mantamadiotis T. Transcriptional regulation of cyclo-oxygenase expression: three pillars of control. Int J Immunopathol Pharmacol. 2003;16:59–67.

30. Barrios-Rodiles M, Tiraloche G, Chadee K. Lipopolysaccharide modulates cyclooxygenase-2 transcriptionally and posttranscriptionally in human macrophages independently from endogenous IL-1 beta and TNF-alpha. Journal of immunology. 1999;163:963–969.

31. D’Acquisto F, Iuvone T, Rombola L, Sautebin L, Di Rosa M, Carnuccio R. Involvement of NF-kappaB in the regulation of cyclooxygenase-2 protein expression in LPS-stimulated J774 macrophages. FEBS letters. 1997;418:175–178.

32. Kang YJ, Wingerd BA, Arakawa T, Smith WL. Cyclooxygenase-2 gene transcription in a macrophage model of inflammation. Journal of immunology. 2006;177:8111–8122.

33. Diaz-Munoz MD, Osma-Garcia IC, Fresno M, Iniguez MA. Involvement of PGE2 and the cAMP signalling pathway in the up-regulation of COX-2 and mPGES-1 expression in LPS-activated macrophages. The Biochemical journal. 2012;443:451–461. doi: 10.1042/BJ20111052

34. Fitzgerald GA. Prostaglandins: modulators of inflammation and cardiovascular risk. Journal of clinical rheumatology : practical reports on rheumatic & musculoskeletal diseases. 2004;10:S12–17. doi: 10.1097/01.rhu.0000130685.73681.8b

35. Liu Y, Wang X, Zhu Y, Cao Y, Wang L, Li F, Zhang Y, Li Y, Zhang Z, Luo J, et al. The CTCF/LncRNA-PACERR complex recruits E1A binding protein p300 to induce pro-tumour macrophages in pancreatic ductal adenocarcinoma via directly regulating PTGS2 expression. Clinical and translational medicine. 2022;12:e654. doi: 10.1002/ctm2.654

36. McAdam BF, Mardini IA, Habib A, Burke A, Lawson JA, Kapoor S, FitzGerald GA. Effect of regulated expression of human cyclooxygenase isoforms on eicosanoid and isoeicosanoid production in inflammation. The Journal of clinical investigation. 2000;105:1473–1482. doi: 10.1172/JCI9523

37. Liu D, Ji L, Wang Y, Zheng L. Cyclooxygenase-2 expression, prostacyclin production and endothelial protection of high-density lipoprotein. Cardiovascular & hematological disorders drug targets. 2012;12:98–105.

38. Norata GD, Callegari E, Inoue H, Catapano AL. HDL3 induces cyclooxygenase-2 expression and prostacyclin release in human endothelial cells via a p38 MAPK/CRE-dependent pathway: effects on COX-2/PGI-synthase coupling. Arteriosclerosis, thrombosis, and vascular biology. 2004;24:871–877. doi: 10.1161/01.ATV.zhq0504.1403

39. Yvan-Charvet L, Wang N, Tall AR. Role of HDL, ABCA1, and ABCG1 transporters in cholesterol efflux and immune responses. Arteriosclerosis, thrombosis, and vascular biology. 2010;30:139–143. doi: 10.1161/ATVBAHA.108.179283

40. Xing Z, Lin C, Yang L. LncRNA Pulldown Combined with Mass Spectrometry to Identify the Novel LncRNA-Associated Proteins. Methods in molecular biology. 2016;1402:1–9. doi: 10.1007/978-1-4939-3378-5_1

41. Feng Y, Chen Z, Xu Y, Han Y, Jia X, Wang Z, Zhang N, Lv W. Corrigendum: The central inflammatory regulator IkappaBzeta: induction regulation and physiological functions. Frontiers in immunology. 2024;15:1398222. doi: 10.3389/fimmu.2024.1398222

42. Trinh DV, Zhu N, Farhang G, Kim BJ, Huxford T. The nuclear I kappaB protein I kappaB zeta specifically binds NF-kappaB p50 homodimers and forms a ternary complex on kappaB DNA. Journal of molecular biology. 2008;379:122–135. doi: 10.1016/j.jmb.2008.03.060

43. Yamazaki S, Muta T, Takeshige K. A novel IkappaB protein, IkappaB-zeta, induced by proinflammatory stimuli, negatively regulates nuclear factor-kappaB in the nuclei. The Journal of biological chemistry. 2001;276:27657–27662. doi: 10.1074/jbc.M103426200

44. Motta-Mena LB, Heyd F, Lynch KW. Context-dependent regulatory mechanism of the splicing factor hnRNP L. Molecular cell. 2010;37:223–234. doi: 10.1016/j.molcel.2009.12.027

45. Rothrock CR, House AE, Lynch KW. HnRNP L represses exon splicing via a regulated exonic splicing silencer. The EMBO journal. 2005;24:2792–2802. doi: 10.1038/sj.emboj.7600745

46. Li Z, Chao TC, Chang KY, Lin N, Patil VS, Shimizu C, Head SR, Burns JC, Rana TM. The long noncoding RNA THRIL regulates TNFα expression through its interaction with hnRNPL. Proceedings of the National Academy of Sciences of the United States of America. 2014;111:1002–1007. doi: 10.1073/pnas.1313768111

47. Atianand MK, Hu W, Satpathy AT, Shen Y, Ricci EP, Alvarez-Dominguez JR, Bhatta A, Schattgen SA, McGowan JD, Blin J, et al. A Long Noncoding RNA lincRNA-EPS Acts as a Transcriptional Brake to Restrain Inflammation. Cell. 2016;165:1672–1685. doi: 10.1016/j.cell.2016.05.075

48. Deng WG, Zhu Y, Wu KK. Role of p300 and PCAF in regulating cyclooxygenase-2 promoter activation by inflammatory mediators. Blood. 2004;103:2135–2142. doi: 10.1182/blood-2003-09-3131

49. Ou J, Tu H, Shan B, Luk A, DeBose-Boyd RA, Bashmakov Y, Goldstein JL, Brown MS. Unsaturated fatty acids inhibit transcription of the sterol regulatory element-binding protein-1c (SREBP-1c) gene by antagonizing ligand-dependent activation of the LXR. Proceedings of the National Academy of Sciences of the United States of America. 2001;98:6027–6032. doi: 10.1073/pnas.111138698

50. De Bosscher K, Vanden Berghe W, Vermeulen L, Plaisance S, Boone E, Haegeman G. Glucocorticoids repress NF-kappaB-driven genes by disturbing the interaction of p65 with the basal transcription machinery, irrespective of coactivator levels in the cell. Proceedings of the National Academy of Sciences of the United States of America. 2000;97:3919–3924. doi: 10.1073/pnas.97.8.3919

51. Kothapalli D, Fuki I, Ali K, Stewart SA, Zhao L, Yahil R, Kwiatkowski D, Hawthorne EA, FitzGerald GA, Phillips MC, et al. Antimitogenic effects of HDL and APOE mediated by Cox-2-dependent IP activation. The Journal of clinical investigation. 2004;113:609–618. doi: 10.1172/JCI19097

52. Mertens A, Verhamme P, Bielicki JK, Phillips MC, Quarck R, Verreth W, Stengel D, Ninio E, Navab M, Mackness B, et al. Increased low-density lipoprotein oxidation and impaired high-density lipoprotein antioxidant defense are associated with increased macrophage homing and atherosclerosis in dyslipidemic obese mice: LCAT gene transfer decreases atherosclerosis. Circulation. 2003;107:1640–1646. doi: 10.1161/01.CIR.0000056523.08033.9F

53. Eren E, Yilmaz N, Aydin O. High Density Lipoprotein and it’s Dysfunction. The open biochemistry journal. 2012;6:78–93. doi: 10.2174/1874091X01206010078

54. Sanson M, Distel E, Fisher EA. HDL induces the expression of the M2 macrophage markers arginase 1 and Fizz-1 in a STAT6-dependent process. PloS one. 2013;8:e74676. doi: 10.1371/journal.pone.0074676

55. Grosser T, Ricciotti E, FitzGerald GA. The Cardiovascular Pharmacology of Nonsteroidal Anti-Inflammatory Drugs. Trends in pharmacological sciences. 2017;38:733–748. doi: 10.1016/j.tips.2017.05.008

56. Vinals M, Martinez-Gonzalez J, Badimon L. Regulatory effects of HDL on smooth muscle cell prostacyclin release. Arteriosclerosis, thrombosis, and vascular biology. 1999;19:2405–2411. doi: 10.1161/01.atv.19.10.2405

57. Shridas P, Bailey WM, Gizard F, Oslund RC, Gelb MH, Bruemmer D, Webb NR. Group X secretory phospholipase A2 negatively regulates ABCA1 and ABCG1 expression and cholesterol efflux in macrophages. Arteriosclerosis, thrombosis, and vascular biology. 2010;30:2014–2021. doi: 10.1161/ATVBAHA.110.210237

58. Hennessy EJ, Sheedy FJ, Santamaria D, Barbacid M, O’Neill LA. Toll-like receptor-4 (TLR4) down-regulates microRNA-107, increasing macrophage adhesion via cyclin-dependent kinase 6. The Journal of biological chemistry. 2011;286:25531–25539. doi: 10.1074/jbc.M111.256206

59. Parhiz H, Shuvaev VV, Pardi N, Khoshnejad M, Kiseleva RY, Brenner JS, Uhler T, Tuyishime S, Mui BL, Tam YK, et al. PECAM-1 directed re-targeting of exogenous mRNA providing two orders of magnitude enhancement of vascular delivery and expression in lungs independent of apolipoprotein E-mediated uptake. Journal of controlled release : official journal of the Controlled Release Society. 2018;291:106–115.

60. Carpenter S, Aiello D, Atianand MK, Ricci EP, Gandhi P, Hall LL, Byron M, Monks B, Henry-Bezy M, Lawrence JB, et al. A long noncoding RNA mediates both activation and repression of immune response genes. Science. 2013;341:789–792. doi: 10.1126/science.1240925

61. Meng H, Sengupta A, Ricciotti E, Mrcela A, Mathew D, Mazaleuskaya LL, Ghosh S, Brooks TG, Turner AP, Schanoski AS, et al. Deep phenotyping of the lipidomic response in COVID-19 and non-COVID-19 sepsis. Clinical and translational medicine. 2023;13:e1440. doi: 10.1002/ctm2.1440

62. Song WL, Lawson JA, Wang M, Zou H, FitzGerald GA. Noninvasive assessment of the role of cyclooxygenases in cardiovascular health: a detailed HPLC/MS/MS method. Methods in enzymology. 2007;433:51–72. doi: 10.1016/S0076-6879(07)33003-6

63. FitzGerald GA, Pedersen AK, Patrono C. Analysis of prostacyclin and thromboxane biosynthesis in cardiovascular disease. Circulation. 1983;67:1174–1177. doi: 10.1161/01.cir.67.6.1174

